# ChimPipe: Accurate detection of fusion genes and transcription-induced chimeras from RNA-seq data

**DOI:** 10.1101/070888

**Authors:** Bernardo Rodríguez-Martín, Emilio Palumbo, Santiago Marco-Sola, Thasso Griebel, Paolo Ribeca, Graciela Alonso, Alberto Rastrojo, Begoña Aguado, Roderic Guigó, Sarah Djebali

## Abstract

**Background:** Chimeric transcripts are commonly defined as transcripts linking two or more different genes in the genome, and can be explained by various biological mechanisms such as genomic rearrangement, read-through or trans-splicing, but also by technical or biological artefacts. Several studies have shown their importance in cancer, cell pluripotency and motility. Many programs have recently been developed to identify chimeras from Illumina RNA-seq data (mostly fusion genes in cancer). However outputs of different programs on the same dataset can be widely inconsistent, and tend to include many false positives. Other issues relate to simulated datasets restricted to fusion genes, real datasets with limited numbers of validated cases, result inconsistencies between simulated and real datasets, and gene rather than junction level assessment.

**Results:** Here we present ChimPipe, a modular and easy-to-use method to reliably identify chimeras from paired-end Illumina RNA-seq data. We have also produced realistic simulated datasets for three different read lengths, and enhanced two gold-standard cancer datasets by associating exact junction points to validated gene fusions. Benchmarking ChimPipe together with four other state-of-the-art tools on this data showed ChimPipe to be the top program at identifying exact junction coordinates for both kinds of datasets, and the one showing the best trade-off between sensitivity and precision. Applied to 106 ENCODE human RNA-seq datasets, ChimPipe identified 137 high confidence chimeras connecting the protein coding sequence of their parent genes. In subsequent experiments, three out of four predicted chimeras, two of which recurrently expressed in a large majority of the samples, could be validated. Cloning and sequencing of the three cases revealed several new chimeric transcript structures, 3 of which with the potential to encode a chimeric protein for which we hypothesized a new role.

**Conclusions:** ChimPipe combines spanning and paired end RNA-seq reads to detect any kind of chimeras, including read-throughs, and shows an excellent trade-off between sensitivity and precision. The chimeras found by ChimPipe can be validated *in-vitro* with high accuracy.

## Background

Chimeric transcripts or chimeras are transcripts whose sequence originates from two or more different genes in the genome [1], and can be explained by several different biological mechanisms at the genomic or the transcriptional level. For its historical relation to cancer, the most well known mechanism is genomic rearrangement. This process brings two genes that are far apart in the germline genome close to each other, and in the same direction, in the cancer genome. The fusion gene thus created can have a deleterious role, either as a protein or as a transcript [1, 2]. Aside from chimeras that are important for their known role in cancer, there are other functional *transcriptional* mechanisms that can also explain the formation of chimeras in normal or tumour cells: polymerase read-through and trans-splicing [1].

As indicated by its name, polymerase read-through occurs when the polymerase reads through one gene into the next, therefore creating a chimera between two adjacent genes. Initially thought to be an exception, this mechanism was found to be widespread in mammals when large collections of ESTs (Expressed Sequence Tags) and cDNAs (complementary DNA) became available and were mapped to the genome [3–5], and when the ENCODE (ENCyclopedia Of DNA Elements) consortium systematically surveyed the transcriptome associated to annotated protein coding genes [6–9]. Read-throughs occur between annotated exons of adjacent genes, preferentially between the penultimate exon of the upstream (5’) gene and the second exon of the downstream (3’) gene [3], resulting in new proteins containing domains from the two parent genes, therefore increasing the diversity of a species proteome [1, 3, 4, 10, 11]. They are also largely conserved across vertebrates [11, 12], and could be a way to regulate the expression of one or both parent genes [12].

Trans-splicing is a splicing mechanism that, unlike the well known cis-splicing, occurs between two different pre-messenger RNA (pre-mRNA) molecules close in the three dimensional (3D) space of the nucleus and thought to belong to the same ‘transcription factory’. If the two pre-mRNAs come from two different genes, a transcriptional chimera is generated [1, 13–16]. The two connected genes can therefore be located distally from each other in the genome, however the chimeric junction must have canonical splice sites. Initially thought to be restricted to trypanosomatidas, this mechanism has gained interest in human research since several studies have found chimeras between genes on different chromosomes or strands, without evidence of underlying genomic rearrangements [13, 14, 16]. One hypothesis is that such trans-spliced transcripts occurring in normal cells would trigger a genomic rearrangement, which will in turn produce a higher quantity of these transcripts (although through a different mechanism), eventually leading to tumorigenesis [13].

But chimeras can also be non-functional, either because they are biological noise from the transcriptional machinery, or because they are technical artefacts from Reverse Transcriptase polymerase chain reaction (RT-PCR) based assays. A biological source of artefactual chimeras is polymerase transcriptional slippage through short homologous sequences (SHS), where the polymerase switches template (or pre-mRNA), in the presence of a short sequence with high similarity to the one it is currently transcribing, in another gene close in the 3D space [17]. This mechanism is reminiscent of the reverse transcriptase (RT) template switching, which can also produce artefactual chimeras in RT-PCR-based experiments [18, 19]. Note that in both cases the chimeric junctions will harbor SHS and non canonical splice sites, however those are not sufficient conditions for a chimera to be an artefact, since RNAse protection assay experiments, which are not RT-PCR-based, have confirmed a number of them [9].

The importance of chimeras lies in their ability to create novel transcripts and proteins, therefore potentially altering the phenotype of cells, individuals or groups of individuals [1, 3, 4, 10, 20]. In the field of cancer, some fusion genes are cancer driver events and can be used as biomarkers or even lead to effective treatment - for instance BCR-ABL1 in chronic myeloid leukemia (CML) [21] or TMPRSS2-ERG in prostate cancer [22, 23]. However not all cancer related chimeras result from genomic rearrangements, since some of them can originate from read-through [24–28], and this mechanism could also be the most prevalent one for certain cancer types, such as CLL [29]. Although chimeras' function have mostly been investigated in relation to cancer, chimeras can also be functionally important in other fields. For instance a chimera produced by trans-splicing, TsRMST, has been shown to interact with pluripotency related transcription factors to control cells' pluripotency [15], and the knock-down of two widely expressed chimeras, CTBS-GNG5 and CTNNBIP1-CLSTN1, in non-neoplastic cell lines, resulted in significant reduction in cell growth and motility [30].

These events were previously detected by RT-PCR-based methods such as EST alignment to the genome [5, 12], or RACEarray followed by RT-PCR, cloning and sequencing [7, 9], however RNA-seq has been shown to be both a more precise and a more sensitive detection method [24]. A growing number of bioinformatic methods have been created to detect chimeras amongst such datasets [31–39].

These programs usually include 3 steps: (1) mapping and filtering for chimeric reads, (2) chimeric junction detection, and (3) chimera assembly and filtering [40]. They rely heavily on an underlying mapper to map the reads to the genome (and optionally to the transcriptome), and make use of two kinds of reads for chimera detection:(1) discordant paired end (PE) reads, i.e. paired end reads where the two mates map in a way that is not consistent with annotated gene structure, e.g. on different chromosomes, and (2) ‘split’ reads, i.e. reads that do not map contiguously to the genome but have to be split or fragmented into several blocks (usually two) to map to the genome ((Figure 1). In addition, the use of one or two kinds of reads for chimeric junction detection allows one to define 3 classes of approaches: (1) the whole paired end approach,(2)the direct fragmentation approach, and (3) the paired end + fragmentation approach[41].

**Figure 1.**
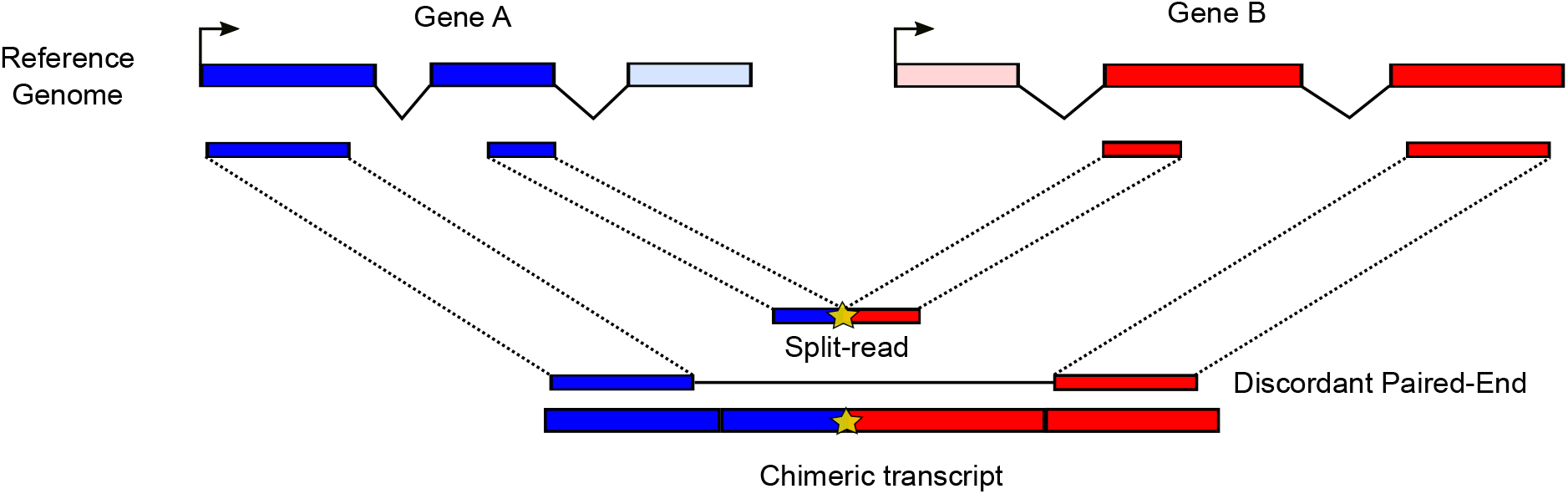
Two types of RNA-seq reads for chimera detection. This picture shows a chimeric transcript (bottom) made from exons of two genes, A and B, depicted in blue and red respectively (bottom). This chimeric transcript is supported by two types of reads: a split-read and a discordant paired-end read, that we depict both on the genome (middle-top) and on the transcriptome (middle-bottom). The chimeric junction position on the transcriptome is highlighted by a yellow star both in the split-read and in the chimeric transcript.

Benchmarking of these programs has shown a high false positive rate and a poor intersection between their outputs on the same dataset [42, 43]. On the other hand these programs are usually developed to detect fusion genes in human cancer, and are therefore not always able to detect read through events and to work on species other than human. In addition, these programs are not always able to predict multiple isoforms per gene pair, and more importantly to provide base pair resolution, preventing their downstream functional validation. To address these problems we present ChimPipe, a modular method which uses the paired end + fragmentation approach and a set of stringent filters, to reliably detect both transcriptional chimeras and fusion genes from Illumina paired-end RNA-seq data from both normal and tumor samples, in any species with a genome and an annotation available. The advantage of the paired end + fragmentation approach is the complementarity of the two types of reads used, with the first ones relatively easy to find but only providing a rough indication of the connected regions, and the second ones more error prone but providing the exact chimeric junction coordinates. ChimPipe represents an advance in methods to quickly and reliably detect chimeric transcripts amongst the rapidly increasing volume of short read transcriptome data.

## Results and discussion

### The ChimPipe method

The ChimPipe method is depicted in Figure 2 and includes 4 consecutive steps:

i. *Exhaustive paired end and split read mapping with GEM*. The paired end reads are initially mapped in 3 ways with the GEMtools RNA-seq pipeline (http://gemtools.github.io/docs/rna_pipeline.html): to the genome, to the transcriptome and *de novo*. Firstly, the reads are mapped to the genome with GEM [44], allowing up to 4% mismatches and indels. Secondly, the reads are mapped to the transcriptome with the same mapping parameters, the transcriptome being composed of all biologically valid combinations of exons within each gene (therefore also including annotated splice junctions). This transcriptome is built from the gene annotation and allows mapping of reads spanning exon to exon junctions that would not match to the reference genome due to the presence of introns. Thirdly, the reads are split-mapped to the genome with the GEM RNA mapper (http://algorithms.cnag.cat/wiki/The_GEM_library) to identify de novo splice junctions from unannotated transcripts. More precisely, reads are split into two segments of at least 15 base pair (bp) length, which are mapped independently to the genome. To reduce the amount of false positive mappings, only split-mappings with less than 4% mismatches or indels and harbouring extended consensus splice sites are further considered (GT+AG, GC+AG, ATATC+A. and `GTATC+AT`, with . meaning any nucleotide). To increase the mapping sensitivity, a second attempt is made by eroding a maximum of two bp towards the ends of each segment if no result is found. At this stage, segments can map to distant positions, but not to different chromosomes, different strands or reverse order. After that, genome, transcriptome and *de novo* mappings are merged and paired and those pairs mapping to more than 10 positions are set as unmapped. Finally, unmapped reads are remapped in a second *de novo* mapping with the GEM RNA mapper. Reads are split-mapped to identify *bona* fide splice junctions connecting loci on different chromosomes, different strands and reverse order. In order to reduce the number of false positive mappings, this last split-mapping is done in a more stringent way since no second attempt is performed if no mapping is found. Note that we chose GEM-based methods for mapping because these programs guarantee that all possible mappings of a read are reported given the input parameters.
ii. *ChimSplice*. Read mapping is followed by candidate chimeric splice junction detection from split-mappings. The split-mapped reads are organized into clusters of reads spanning the same splice junction. The donor and acceptor splice sites are considered when building the clusters to guarantee that all of them are in the 5’ to 3’ orientation. This is very important to determine which are the upstream and downstream parent genes, and is particularly useful in case of unstranded RNA-seq data. Once the clusters have been generated, *ChimSplice* produces a consensus splice junction defined by the exact junction coordinates, the upstream coordinates of the upstream cluster, and the downstream coordinates of the downstream cluster. Additionally, each consensus junction is associated with the number of supporting split-reads and *staggered split*-reads. The term staggered split-reads refers to those reads spanning the same junction but mapping to different external positions and, as a consequence, producing a characteristic ladder-like pattern of reads across the junction (see Figure 2B). This pattern has been suggested to be specific to genuine chimeric transcripts, while false positives usually lack it [45]. This information is recorded and can be used to distinguish real from artefactual chimeras in the downstream *ChimFilter filtering* module. Then, the consensus junctions are annotated. Each junction is compared to the annotated exons in order to determine its two parent genes. In case a junction side overlaps several exons from different genes, the one with a higher overlap is selected. Finally, splice junctions connecting exons from two different genes (chimeric junctions) are selected for downstream analyses.
iii. *ChimPE*. Once chimeric junction candidates have been found using *ChimSplice, ChimPE* looks for further paired end support for them (Figure 2B). Genome, transcriptome and *de novo* mappings are filtered to select only those PE reads with both mates mapped. Those PE reads are compared to annotated exons in the same way as described in (ii), and reads with both mates mapping to exons from different genes are identified (discordant PE reads). For each chimeric junction, discordant PE reads connecting their parent genes are then selected and their relative mapping position to the chimeric junction is evaluated. This is done in order to know whether the discordant PE reads support the existence of the chimeric junction (consistent PE) or if, on the other hand, they are incompatible with the chimeric junction (inconsistent PE). Inconsistent PE can be due to different reasons: they may come from a different chimeric RNA isoform than the one highlighted by *ChimSplice*, or from PE read misalignment, but they could also indicate a *ChimSplice* false positive. Finally, each chimeric junction is associated to the number of consistent and inconsistent PE reads, which can be used in the downstream *ChimFilter* filtering module to filter out artefactual chimeras.
iv. *ChimFilter.*Chimeric junction candidates are filtered to produce a final set of more reliable chimeras. Firstly, based on the principle that false positives due to read misalignment would not be supported by both sources of evidence, ChimPipe requires a candidate chimera to be supported by both split-reads and consistent PE reads. Two different support based filtering schemes are applied depending on whether the chimeric junction involves annotated or novel splice sites. By default, chimeric junctions with annotated splice sites must be supported by at least one consistent PE read, one split-read and three total (consistent PE + split) reads, while those with novel splice sites have to be supported by at least three consistent PE reads, three split-reads and six total reads. Secondly, chimeras between genes that share high exonic sequence similarity are less reliable since the supporting reads are more prone to mis-alignments. For this reason ChimPipe implements an homology-based filtering and chimeric junctions connecting genes with homologous exonic sequence of at least 30 bp (and 90% sequence identity), are discarded as likely false positives. All these filtering parameters can be tuned.

The main ChimPipe output is a tabulated text file with header including the set of chimeric junctions after filtering, in which the first column is the junction identifier in ChimPipe format (donchr_donpos_donstr:accchr_accpos_accstr), and the other 34 columns are valuable pieces of information about it, such as its support in terms of number of staggered split and consistent discordant paired-end reads, its type *(readthrough* (resp. *intrachromosomal*) if the two parts are on the same chromosome, same strand, expected genomic order and closer (resp. more distant) than 100 kilobase (kb), *inverted* if the two parts are on the same chromosome, same strand and unexpected genomic order, *interstrand* if the two parts are on the same chromosome but different strands and *interchromosomal* if the two parts are on different chromosomes), its two parent genes, its length and the list of its supporting reads (see Additional data file 1 for more details). ChimPipe also outputs a file with chimeric junctions before the filtering, and a file with the junctions that have been filtered out with information about the reason for this filtering (see ChimPipe user’s manual at http://chimpipe.readthedocs.io/en/latest/manual.html for more information). It has to be noted that ChimPipe can also start from already aligned reads (bam file) provided they include evidence of intra-chromosomal chimeric junctions, and that ChimPipe does not only output chimeric junctions but also a standard bam file (from step (i) of the pipeline) that can be used for more standard RNA-seq analyses such as differential gene expression or transcript reconstruction. Finally ChimPipe has been designed to require minimal information about the PE RNA-seq dataset on which it is run, since it guesses the Illumina quality offset, the strandedness, and the mate configuration in case of directional data. Note that ChimPipe’s documentation includes a tutorial and an example.

**Figure 2.**
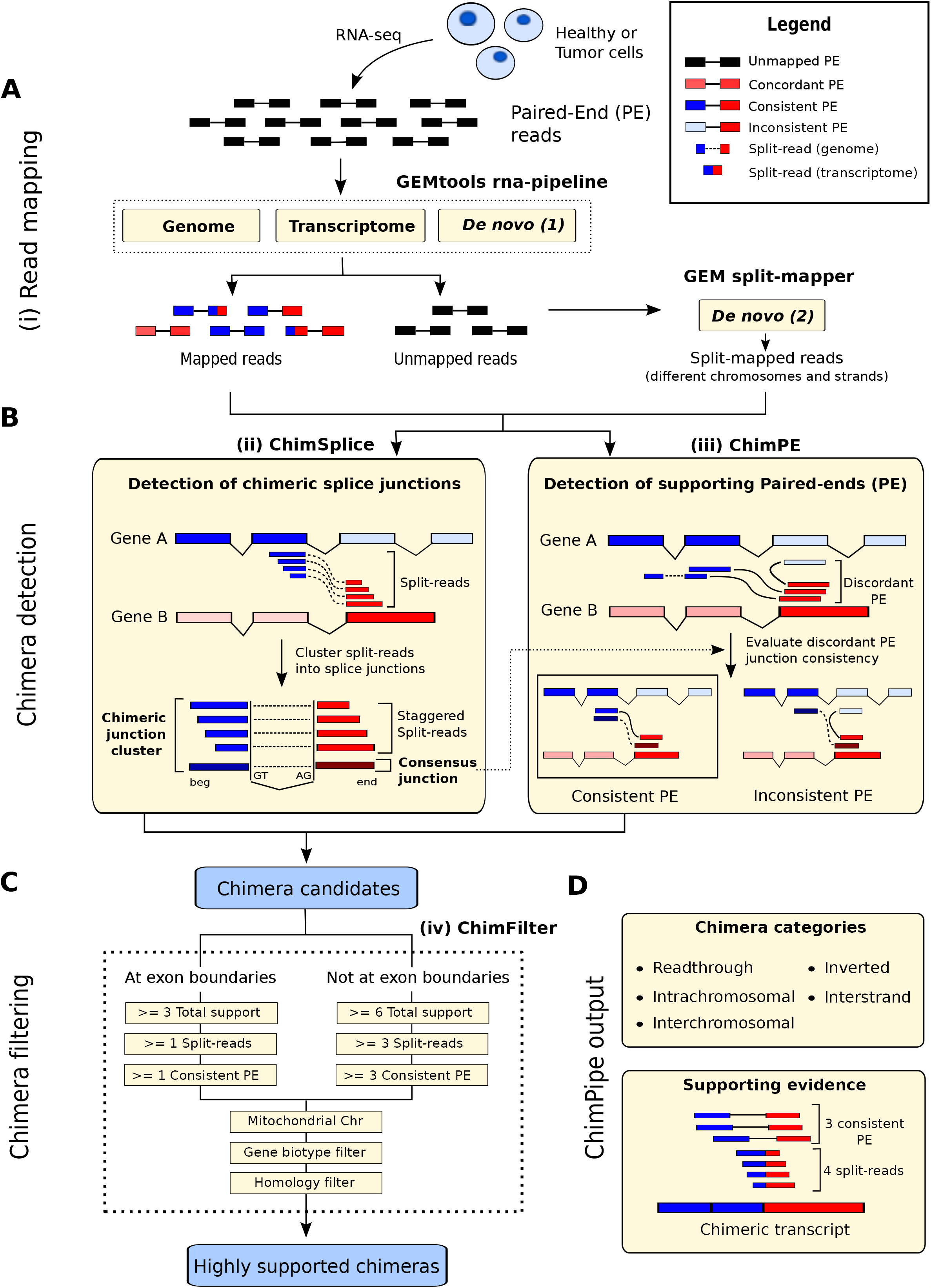
The ChimPipe method. (A) RNA-seq reads are first mapped to the genome and transcriptome using the GEMtools RNA-seq pipeline, and the reads that do not map this way are passed to the GEM RNA-mapper to get reads that split map to different chromosomes or strands. (B) The split-reads from these two mapping steps are then gathered and passed on to the *ChimSplice* module which derives consensus junctions associated to their expression calculated as the number of staggered split-reads supporting them. The *ChimPE* module can then associate each chimeric junction found by *ChimSplice* to their discordant PE reads, splitting them into the ones consistent and the ones inconsistent with the junction. (C) The *ChimFilter* module then applies a series of filters to the chimeric junctions obtained until this point in order to discard false positives, leading to (D) a set of reliable chimeric junctions to which it associates several pieces of information such as a category (readthrough, intrachromosomal, inverted, interstand, or interchromosomal), and the supporting evidence in terms of number of staggered split-reads and number of consistent PE reads, among others.

### Benchmark on simulated and cancer data

We evaluated ChimPipe and other state-of-the-art chimera detection tools, using two kinds of evaluation data: simulated data and real data from melanoma and breast cancer. The main advantages of simulated data are the inclusion of all kinds of chimeras (not only fusion genes) and the control over the chimeras expected to be found, therefore allowing a precise evaluation of the programs. Its main drawback, however, is the uncertainty about whether it captures the underlying complexity of real data. The drawback of real data, on the other hand, is its very limited number of validated cases, and the fact that most of them are fusion genes. Indeed neither does it allow to assess the programs’ precision, nor to extrapolate their results to non cancer data.

We developed `ChimSim`, a program to simulate chimeric transcripts from a gene annotation, a genome, and numbers of read-through, intra-chromosomal, inverted, interstrand and interchromosomal chimeric transcripts to create from the annotation (see supplementary methods). Using `ChimSim` on the the Gencode v19 protein-coding genes [46] and the hg19 genome, we generated a simulated dataset of 250 chimeric transcripts homogeneously distributed in the 5 chimera classes (50 from each class) (Additional data file 2). Knowing that about 60% of transcripts from protein coding (pc) and long non-coding RNA (lncRNA) genes are usually expressed in a given condition [47], we sampled 60% of transcripts from Gencode v19 pc and lncRNA gene transcripts, totalling 101,961 transcripts (Additional data file 2). Knowing that when a chimera is expressed, its parent genes are often also expressed [10], we added the parent transcripts of the 250 chimeras to the sampled transcripts, totalling 102,149 non chimeric transcripts (Additional data file 2).

The 102,399 transcripts resulting from the union of the 250 chimeric transcripts and the 102,149 non-chimeric transcripts, were then passed on to the `art_illumina` program of the ART suite (version 2.3.7, [48]), to simulate Illumina non directional paired-end RNA-seq reads of 3 different lengths: 50bp, 76bp and 101bp, called PE50, PE76 and PE101 respectively. Several parameters were used in addition to read length and paired-endness, to make our simulated chimera data closer to real RNA-seq data, including insert size mean and standard deviation, read coverage and sequencing quality profile (see supplementary methods for details). The sequencing quality profile was learnt from real Illumina PE data of the same read length using the `art_profiler_illumina` program of the ART suite (version 2.3.7, Additional data file 2 and supplementary methods). Using these parameters, ART generated 32.3, 21.1 and 15.7 million PE reads for the PE50, PE76 and PE101 respectively (Additional data file 2 and supplementary methods). The benchmark was done for each read length separately.

For real data with experimentally validated chimeras, we used two previously published datasets: the leukemia/melanoma cancer study from Berger et al ([25], that we call the `Berger set`), and the breast cancer study from Edgren et al ([45], that we call the Edgren set). The Berger set was composed of the K562 chronic myelogenous leukemia cell line associated to two different insert size ranges, 300-400 bp and 400-600 bp, of the 501Mel melanoma cell line and of 5 melanoma patient-derived short-term cultures, and came with 14 RT-PCR validated fusion genes (Table 1). The Edgren set was composed of 4 breast cancer cell lines (of which two were associated to two different median insert sizes, 100bp and 200bp), and came with 27 RT-PCR validated fusion genes. For the Edgren set we used an additional 13 fusion genes that were found and RT-PCR-validated by a re-analysis of the Edgren data by Kangaspeska et al. [49], totalling 40 fusion genes (Table 1). The benchmark was done for each library separately, but is provided for the pool of libraries of each dataset, for clarity reasons. Since the chimeras specifically targeted by the Berger and the Edgren studies were only fusion genes, the read-through events were removed from all programs’ predictions before running the benchmark.

**Table 1.** This table indicates for each cancer dataset, its associated set of cell lines and corresponding tumor types, together with the number of RT-PCR validated fusion genes and junctions. Some fusion genes are associated to several fusion junctions.

| Cancer dataset | Cell line | Tumor type | Number of validated fusion genes | Number of validated fusion junctions | Number of different libraries (insert sizes) | SRA* accession codes |
| --- | --- | --- | --- | --- | --- | --- |
| Berger | K-562 | Leukemia | 3 | 3 | 2 | SRR018268, SRR0182689 |
|  | 501 Mel | Melanoma | 4 | 5 | 1 | SRR018266 |
|  | M000216 |  | 1 | 1 | 1 | SRR018259 |
|  | M000921 |  | 2 | 3 | 1 | SRR018267 |
|  | M010403 |  | 1 | 1 | 1 | SRR018265 |
|  | M980409 |  | 1 | 1 | 1 | SRR018261 |
|  | M990802 |  | 2 | 2 | 1 | SRR018260 |
|  | All | All | **14** | **16** | - | - |
| Edgren | KPL-4 | Breast cancer | 3 | 3 | 1 | SRR064287 |
|  | MCF-7 |  | 6 | 8 | 1 | SRR064286 |
|  | BT-474 |  | 21 | 25 | 2 | SRR064438,SRR064439 |
|  | SK-BR-3 |  | 10 | 10 | 2 | SRR064440,SRR064441 |
|  | All |  | **40** | **46** | 1 | - |

\*SRA: Sequence Read Archive (http://www.ncbi.nlm.nih.gov/sra)

Since we wanted to do the evaluation both at the gene pair level and at the chimeric junction level, and since an RT-PCR validated fusion gene is merely a gene pair together with the cDNA sequence corresponding to its junction, we used the blat program [50] to align the cDNA sequences to the hg19 human genome, and further manually curated these alignments to obtain the exact chimeric junction coordinates for each fusion gene (see supplementary methods). This procedure resulted in 16 and 42 chimeric junctions for the Berger and Edgren sets respectively, indicating the presence of two different isoforms for one gene pair in each set (Table S1).

The chimera detection programs that we chose to benchmark together with ChimPipe (version 0.9.3) were the following:

- FusionMap (version 8.0.2.32, [33])
- PRADA (version 1.2, [38])
- Chimerascan (version 0.4.5, [34])
- TopHatFusion (version 2.0.12, [32]).

We chose these programs because their method was published and for one of the following three reasons: (1) they were shown to have good results in several independent studies (for example FusionMap and Chimerascan) or (2) they were used in studies associated with gold-standard chimera RNA-seq datasets (for example PRADA in [25] and Chimerascan in [24]) or (3) they were extensively used by the community (for example TopHatFusion). Since these programs are optimized for human and for cancer, we used them with their default parameters for real data, and we adjusted their parameters to allow read-through detection for simulated data, whenever it was possible (see supplementary methods).

The evaluation measures used are the standard sensitivity and precision for simulated data, and the sensitivity and total predictions for real data. In addition, the evaluation was done at two levels: the gene pair level and the junction level (Figure S1).For each of these two objects, gene pair and junction, we have a reference set (the objects to be predicted), and a predicted set for each program (the objects actually predicted by the program). We then define a true positive (TP) as an object present both in the reference and in the predicted set, a false positive (FP) as an object present in the predicted set but not in the reference set, and a false negative (FN) as an object present in the reference set but not in the predicted set. The sensitivity (Sn) is then the fraction of the reference objects that are correctly predicted, while the precision (Pr) is the fraction of the predicted objects that are correctly predicted. Since a high Sn can be easily obtained at the expense of a low Pr, and reciprocally, we use the average of Sn and Pr as an additional measure. Note that in order for a predicted chimeric junction to be a TP, its coordinates must exactly match the coordinates of a reference chimeric junction (Figure S1 and supplementary methods).

The results at both the gene pair level and at the junction level for both the PE76 simulated data and the real data, are shown on Figure 3 (similar results were observed for PE50 and PE101 except for FusionMap which is clearly better on PE76, see Figure S2). At the gene pair level the top program on the simulated data is Chimerascan followed by ChimPipe, FusionMap, PRADA and finally TopHatFusion, with a generally quite high precision for all programs but a sensitivity above 0.75 only for Chimerascan and ChimPipe. For real data, Chimerascan is still the top program in terms of sensitivity followed by ChimPipe, however its number of predicted gene pairs is 1 to 2 orders of magnitude higher than the one of ChimPipe. The trend for real data sensitivity is similar to the one of simulated data, but the Edgren gene pairs seem to be easier to predict than the Berger gene pairs, with a higher sensitivity of the programs for the former. Note that PRADA is a program that also has a good compromise between Sn and number of predicted gene pairs on real data.

**Figure 3.**
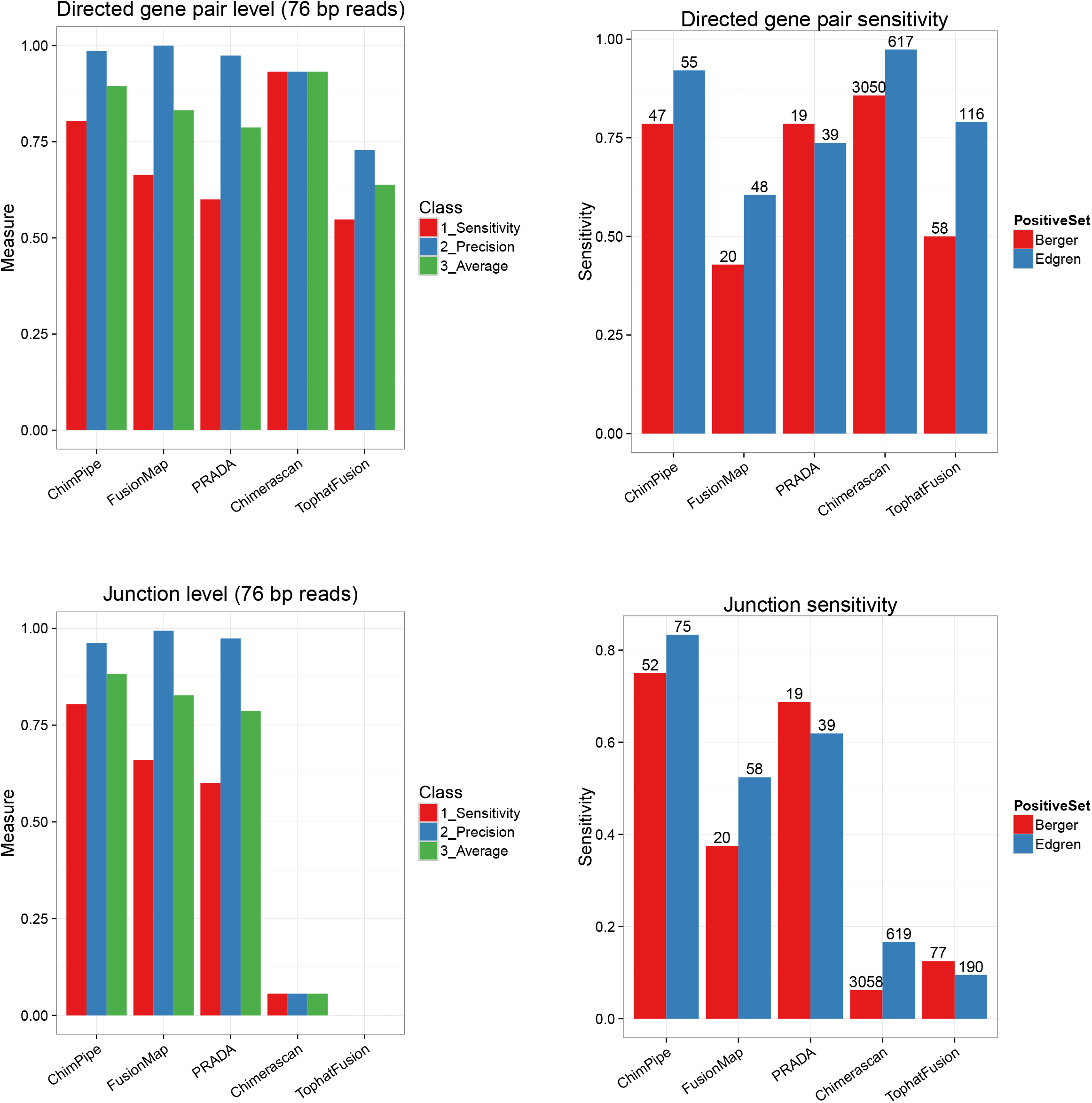
Benchmark results for 5 chimera detection programs on simulated (left) and on real (right) data. The sets of barplots on the top (A,B) indicate the programs’ performances at the gene pair level, while the sets of barplots at the bottom (C,D) indicate the programs’ performances at the junction level. For simulated data the provided measures are sensitivity (in red), precision (in blue), and average between the two (in green), while for the two real datasets (Berger in red and Edgren in blue), the only provided measures are sensitivity (bars) and the total number of predictions (at the top of each bar). Here we show the results on PE76 simulated data, for the 250 simulated chimeric junctions (i.e. including read through events). For the benchmark on real data, read through events, i.e. junction distance smaller than 100kb when on the same chromosome, same strand and expected genomic order, were removed from the output of each program before the evaluation.

At the junction level, ChimPipe is the best program on both the simulated and the real data with a sensitivity around 0.8 and a precision close to 1, and with a quite reasonable number of predicted junctions for real data (around 60). It is followed by PRADA and FusionMap, with PRADA behaving clearly better on real data. The performances of both Chimerascan and TopHatFusion are quite poor at the junction level, with TopHatFusion junctions always shifted by 1 bp on each side, and Chimerascan junctions most often shifted by 1 bp on one or both sides, compared to true junctions.

Since some of the evaluated programs are not able to predict read-through events (PRADA), or happened to not detect any of them on simulated data (FusionMap and TopHatFusion), we also made an evaluation without read-through events on simulated data (Figure S3). The effect was an overall improvement of the programs’ performances (except ChimPipe) but did not change the overall message above.

Since some programs have a quite different behaviour at the gene pair and at the junction level, we also computed for each program and each evaluation set, the average and standard deviation of the distance between the predicted and the true junction in case the gene pair was correctly predicted (Table 2). It showed that ChimPipe, FusionMap and PRADA almost always provide the exact junction coordinates on simulated data and the Berger real set, while this is not the case for Chimerascan and TopHatFusion, with a worse behaviour for the latter on the simulated data and for the former on the Berger real set. One can note that for simulated data, the distance between Chimerascan predicted and true junction tends to increase with read length. Although the Edgren gene pairs seem easier to predict than the Berger gene pairs (as stated above), the junctions from the correctly predicted gene pairs seem more difficult to predict for the Edgren set than for the Berger set, since all the programs show a quite important average distance between predicted and true junction for the Edgren set. ChimPipe is second after FusionMap on the Edgren set but also has many more true positives on this set. Since when ChimPipe detects the correct gene pair it also detects the correct junction both for the simulated data and for the Berger cancer data, we think that the most likely explanation for this difficulty in finding the true junction for some Edgren cases is the fact that the mRNA isoform represented by the RT-PCR sequence is not the same as the one sequenced with RNA-seq.

**Table 2.**
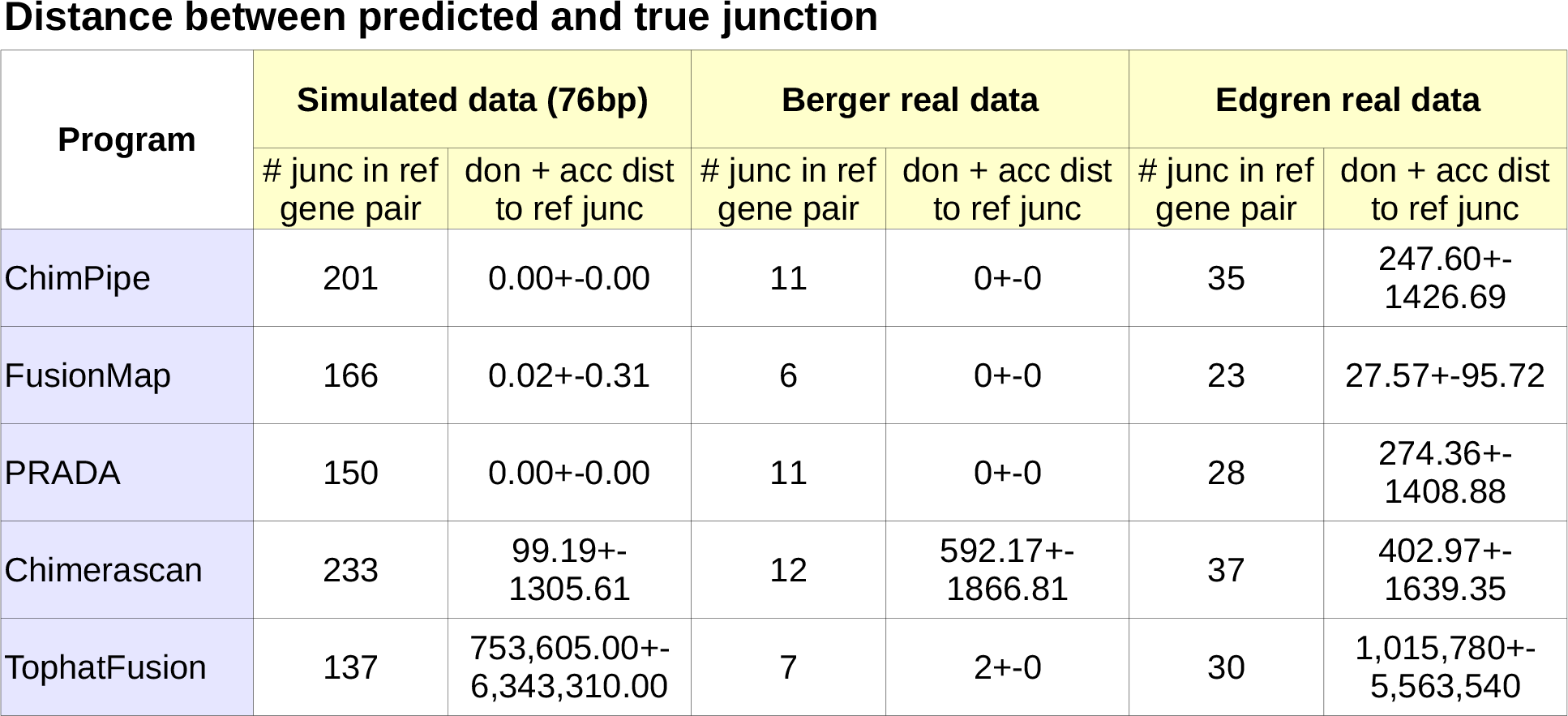
This table provides the number of predicted chimeric junctions corresponding to a reference gene pair and the average and standard deviation of the distance between the two (calculated as the sum of the donor and the acceptor distances), for each program and each benchmark dataset

Although real data does not allow the computation of precision or false positive rate, we expect the number of programs predicting a given chimera to be correlated to the likelihood of this chimera being a true positive. We computed the intersection between the gene pairs predicted by each program on each of the two real sets (Berger and Edgren) (Figure 4), and saw that PRADA, ChimPipe and FusionMap predicted fewer unique gene pairs, while TopHatFusion and Chimerascan predicted many unique gene pairs, consistent with the previous benchmark results (Figure 3). We also confirmed that a gene pair predicted by at least 2 programs was more likely to be real since 26% (respectively 65%) of the gene pairs predicted by 2 programs on the Berger (resp. Edgren) set are true positives (i.e. validated by RT-PCR), while only 0% (respectively 1%) of the ones predicted by 1 program are true positives.

**Figure 4.**
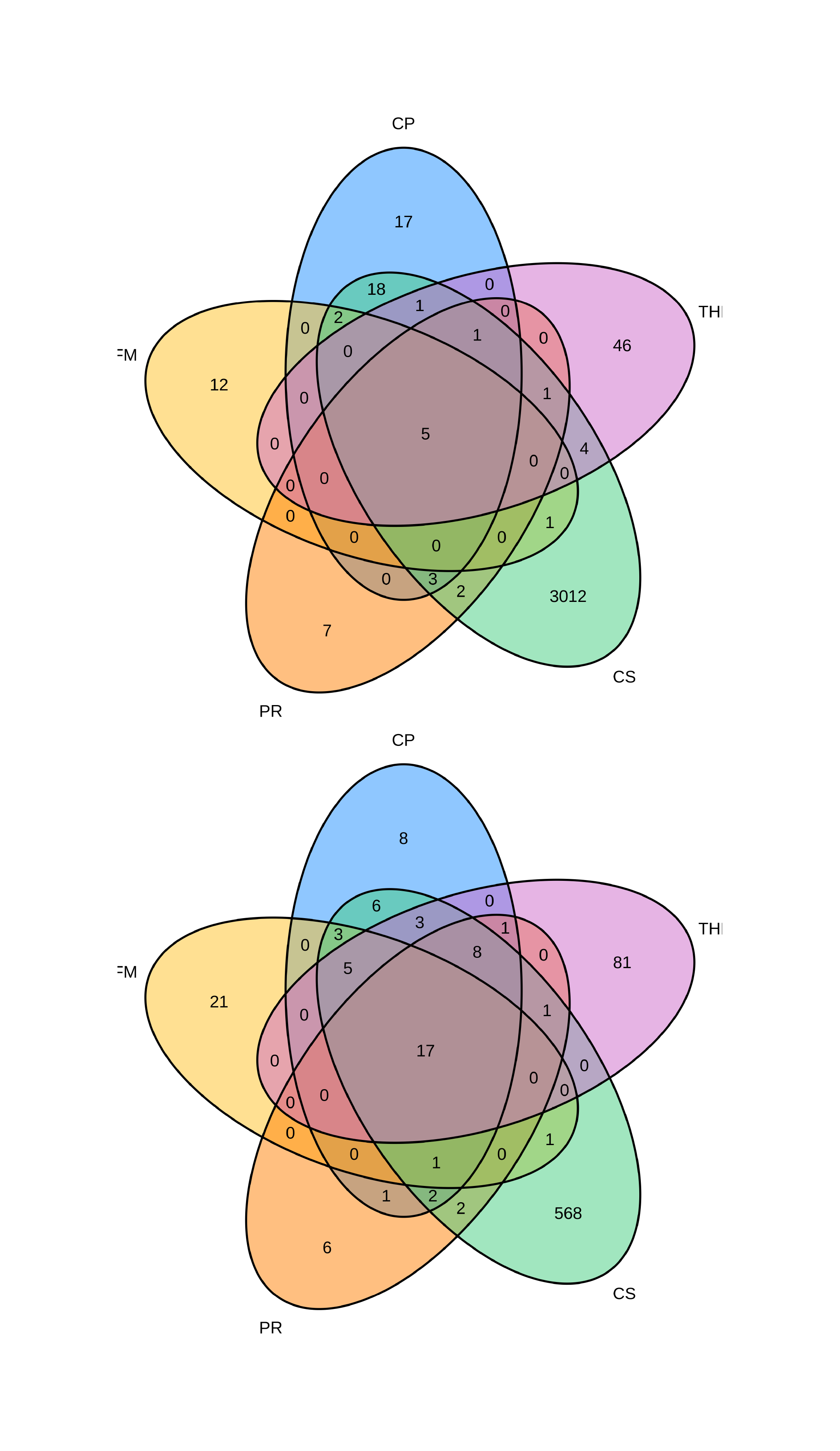
Chimeric gene pairs predicted by the 5 programs on the two real datasets. Intersection between chimeric gene pairs predicted by the 5 programs on the Berger set (A) and on the Edgren set (B) are represented as Venn diagrams. In general gene pairs predicted by all 5 programs are few compared to the gene pairs predicted by a single program, and we expect that the higher the number of programs predicting a gene pair the more reliable the gene pair. Chimerascan and TophaFusion are the programs that predict more gene pairs predicted by no other program, while PRADA, Chimpipe and FusionMap are the programs with less such gene pairs. CP: ChimPipe, FM: FusionMap, PR: PRADA, CS: Chimerascan, THF: TopHatFusion.

Regarding implementation, while some programs require a single step (apart from the genome and/or transcriptome indexing) to predict the chimeras, which is the case for ChimPipe, FusionMap and Chimerascan, some other programs require many different successive steps to obtain them, making the whole process more cumbersome. This is the case for PRADA which requires 3 steps (mapping script making + mapping + chimera prediction) and for TopHatFusion which requires 2 steps (mapping + mapping filtering). The maximum virtual memory and wallclock time needed by each program (run with 4 threads) on the PE76 simulated data are provided in Table 3. The program that clearly needs the least resources is FusionMap with 11.7 Gb of RAM and less than half an hour of running time, followed by Chimerascan with 4.8 Gb of RAM and 8.2 hours of running time, then PRADA with 35.5 Gb of RAM and 4.5 hours of running time, then ChimPipe with 34.5 Gb of RAM and 10.1 hours of running time, and finally TopHatFusion which requires 62.2 Gb of RAM and 18 hours of running time.

**Table 3.** This table indicates the computing resources needed by each program to process the PE76 simulated data, as well as the number of commands needed to produce the final result

| Program | Maximum RAM used (in Gb) | Cumulative wallclock time (in hours) | Number of commands to execute |
| --- | --- | --- | --- |
| ChimPipe | 34.5 | 10.1 | 1 |
| FusionMap | 11.7 | 0.4 | 1 |
| PRADA | 35.5 | 4.5 | 3 (make mapping script, mapping, compute fusion) |
| Chimerascan | 4.8 | 8.2 | 1 |
| TophatFusion | 62.2 | 18* | 2 (mapping, filtering) |

\* 9 hours for mapping and ~9 hours for filtering (27.5 hours for the 3 simulated sets (50bp, 76bp and 101bp) at the same time)

### Detection and validation of novel chimeras

In order to survey the human chimera landscape more extensively, ChimPipe was run on 106 ENCODE RNA-seq experiments from 15 human cell lines, 3 RNA fractions (polyadenylated, non-polyadenylated, total) and 6 cell compartments (whole cell, cytosol, nucleus, chromatin, nucleolus, nucleoplasm) [47]. At stringent settings (10 supporting staggered split-reads and 5 discordant paired end reads in at least one experiment), we found a total of 1195 chimeric junctions over all experiments. Of these, 525 had their 5’ and 3’ ends falling in a single protein-coding gene respectively, and 142 were either expressed recurrently (at least 1 supporting read in at least 11 out of the 15 cell lines) or very highly and specifically (at least 100 total reads in a single cell line). We then only considered the 137 read-through and intrachromosomal chimeric junctions from this set (Additional data file 3).

Four of these chimeric junctions were chosen for RT-PCR plus Sanger sequencing validation. Two of them were selected from the recurrently expressed class (RPL38-TTYH2 and UBA2-WTIP), and two of them from the very highly and specifically expressed class (PICALM-SYTL2 and C16orf62-IQCK) (Table 4). Primers were designed to perform RT-PCR on cDNA (to test for the RNA chimera) as well as PCR on genomic DNA, to assess whether the chimeras could originate from genomic rearrangements (Figure S4, Additional data file 3). Out of those 4 cases, all showed evidence of the two parent gene mRNAs (except one, SYTL2, but this could be due to a low expression level of this gene), and 3 showed the additional presence of the chimeric RNA (Figure S5 and supplementary methods). These 3 chimeric junctions present at the RNA level, were not present at the DNA level and therefore cannot originate from genomic rearrangements (Figure S6 and supplementary methods). We cloned and sequenced these three chimeras (UBA2-WTIP, PICALM-SYTL2 and RPL38-TTYH2) (Figure S7, Figure 5B for UBA2-WTIP, Additional data file 3 and supplementary methods). These data are consistent with the hypothesis that some chimeras originate from read-through or trans-splicing events.

**Figure 5.**
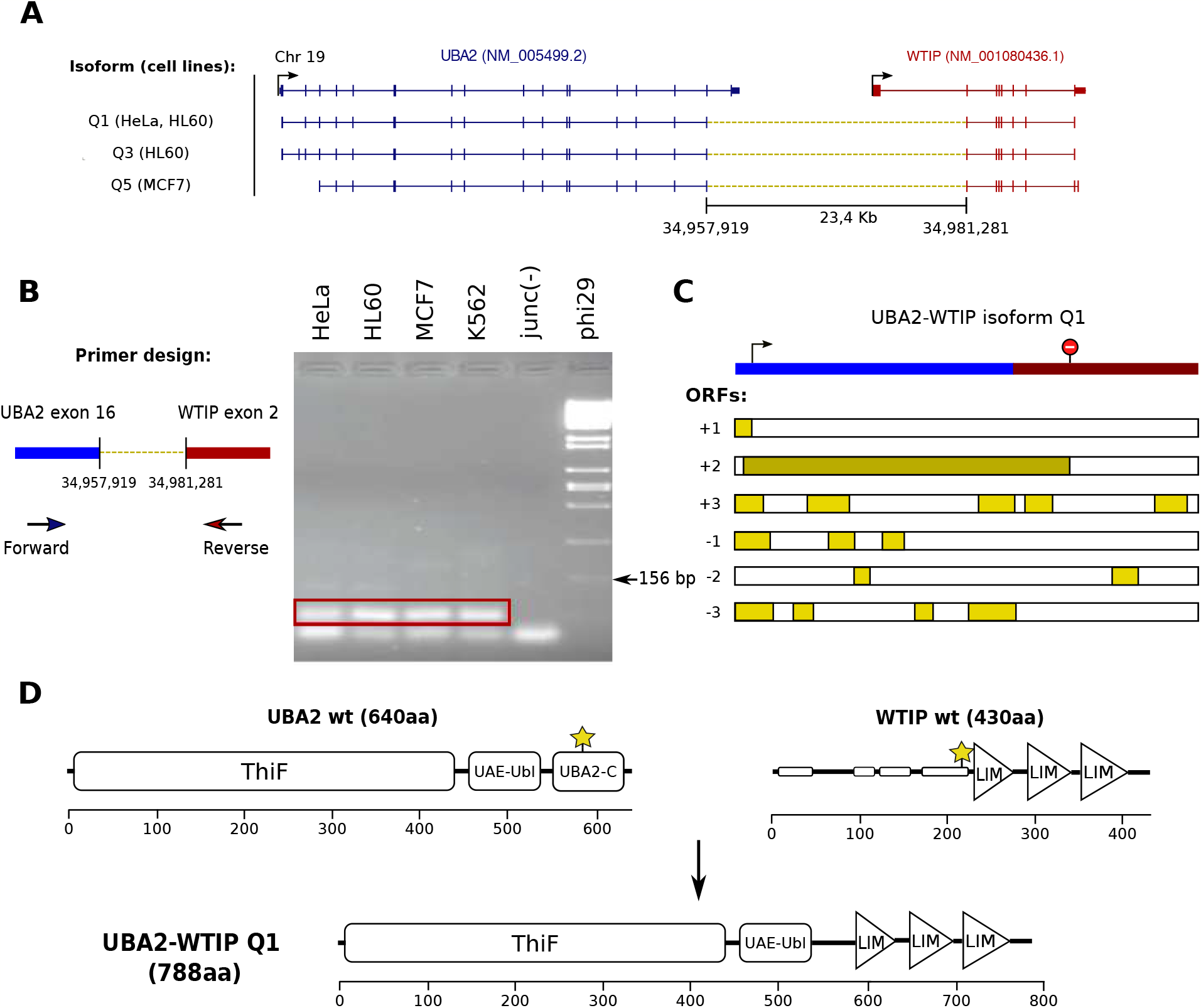
UBA2-WTIP chimeric transcript isoforms. (A) Experimentally validated UBA2-WTIP chimeric transcript isoforms. (Top) UBA2 and WTIP parent transcripts according to RefSeq version 74. Coding and UTR exonic sequences are displayed as thick and thin boxes, respectively, and introns as lines. The genomic strand of the transcripts is represented as an arrow on the 5’ end (Bottom) Chimeric RNAs with chimeric splice junctions are depicted as yellow dashed lines. On the left, list of cancer cell lines where each isoform was validated (B) UBA2-WTIP chimeric splice junction validation (Left) Primer design for validating the chimeric junction through RT-PCR plus Sanger sequencing. (Right) Chimeric junction validation in 4 different cell lines. The 72 bp amplicons proving the expression of the chimeric RNAs are highlighted in red. (C) UBA2-WTIP Q1 isoform protein coding potential. (Top) UBA2 and WTIP annotated start and stop codons represented over the transcript sequence. (Bottom) ORFs in the six possible frames. The selected ORF from the UBA2 annotated start codon to the WTIP annotated stop codon is highlighted in dark yellow. (D) Putative chimeric protein encoded by the UBA2-WTIP Q1 isoform. (Top) UBA2 and WTIP wild type proteins. The exact position of the two protein breakpoints is indicated by yellow stars. Protein domains are depicted as boxes and triangles over the protein sequences. Thin boxes on the WTIP protein sequence correspond to low complexity regions. The x axis shows the amino acid position along the protein sequence. (Bottom) Putative UBA2-WTIP chimeric protein.

**Table 4.**
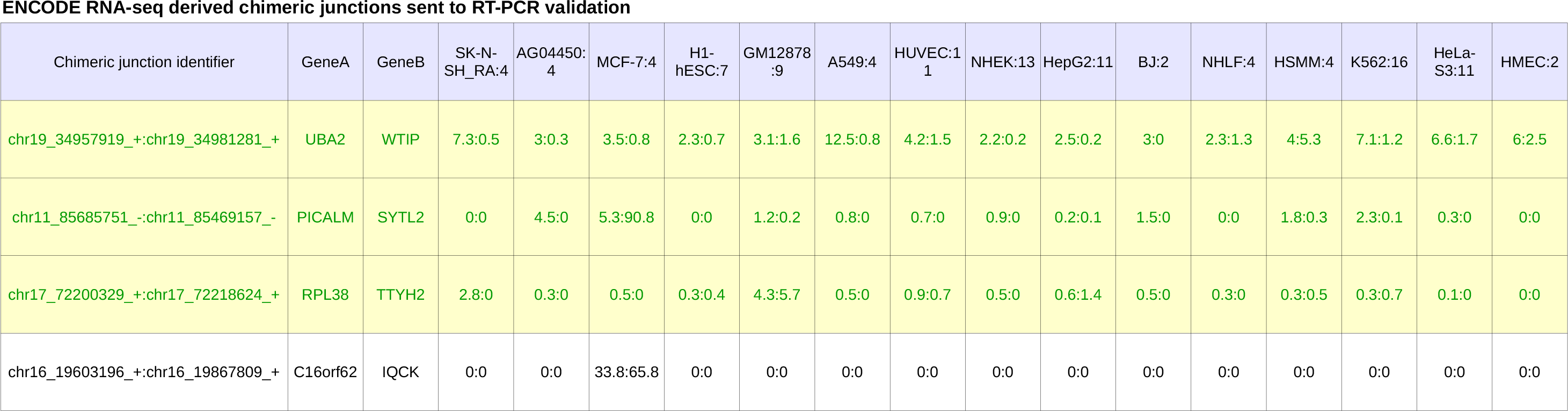
This table lists the 4 chimeric junctions derived from the 106 ENCODE RNA-seq experiment, that we selected for RT-PCR validation (the successful ones are indicated in green on a yellow background). For each junction, its identifier (junction coordinates in ChimPipe format), the name of its 5’ and 3’ genes, and its expression in terms of number of staggered split-reads and discordant paired-end reads in each of the 15 ENCODE cell lines (separated by “:”), are provided. Since each cell line is associated to several RNA-seq experiments (number indicated in the header after the “:” sign), average number of staggered split-reads and of discordant paired end reads across experiments of a cell line are provided.

It has been suggested that the generation of chimeric transcripts and their translation into chimeric proteins may serve to generate novel proteins with altered functions [1, 10]. Therefore, we assessed the protein-coding potential of the 3 validated chimeric junctions. For each chimeric junction, we reconstructed the theoretical chimeric transcript structures by combining the RefSeq reference mRNAs for the 5’ and 3’ parent genes compatible with the junction and searched for Open Reading Frames (ORFs) in the six possible translational frames with the NCBI ORF Finder (http://www.ncbi.nlm.nih.gov/gorf/gorf.html). One case out of the three (UBA2-WTIP), maintained the frame of the two parent genes, UBA2 and WTIP, while the other two, PICALM-SYTL2 and RPL38-TTYH2, did not. Interestingly, this chimera is recurrently expressed in 72 out of the 106 experiments, which include the 15 cell lines, the 3 RNA fractions and 5 out of the 6 cell compartments (cell, cytosol, nucleus, nucleoplasm and chromatin) (Additional data file 3). Additional RT-PCR and Sanger sequencing was therefore performed on UBA2-WTIP, giving rise to 3 novel complete transcript structures (Figure 5A, Additional data file 3), of which the longest one (Q1), was more deeply analysed here. This complete chimeric transcript has an ORF from UBA2 to WTIP annotated start and stop codon respectively (Figure 5C). Thus, if translated it would give rise to a chimeric protein including the two most N-terminal domains of the 5’ parent protein UBA2 (ThiF and UAE_Ubl domains) and the three most C-terminal domains of the 3’ parent protein WTIP (LIM domains), therefore only skipping the UBA2_C domain of the UBA2 protein and the proline-rich N-terminal domain of the WTIP protein (Figure 5D).

We further investigated the putative role of this chimeric protein containing the combination of domains from UBA2 and WTIP wild type proteins. UBA2 is part of the SUMOylation machinery, which post-translationally modifies and regulates a large number of proteins with important roles in diverse cellular processes, including regulation of transcription, chromatin structure, and DNA repair [51]. More precisely, it associates with the Aos1 protein to produce the SUMO-activating enzyme (E1), a heterodimer that mediates the activation of ubiquitin-related modifier (SUMO) molecules and their transference to the SUMO-conjugating enzyme (E2), which post-translationally modifies a target protein through the binding of SUMO [52]. On the other hand, WTIP belongs to a subset of LIM-domain containing proteins, which are involved in focal and cell-cell adhesion. These interact with other proteins through their LIM domains, whose sequence specifies a double zinc-finger structure capable of high-affinity binding to a wide variety of protein targets [53]. Based on this, we hypothesize that the combination of UBA2 SUMOylation domain and WTIP protein binding LIM domains could lead to a chimeric protein with altered SUMOylation activity. This protein may induce the SUMOylation machinery to post-translationally modify and regulate novel targets, due to the interaction of its LIM-domains with novel proteins.

Finally, each one of the two other validated chimeras, PICALM-SYTL2 and RPL38-TTYH2, gave rise to one novel (although incompletely identified) transcript structure with a premature stop codon before the last splice junction, leading us to hypothesize that they are degraded through nonsense-mediated mRNA decay [54]. However, it is important to note that these chimeric junctions are supported by a very high number of reads (Table 4), suggesting that the chimeric transcripts is highly expressed, and possibly functional.

## Conclusions

We have presented ChimPipe, a novel method for the accurate detection of chimeras from PE RNA-seq data, based on the independent use of discordant PE reads and split reads. In addition to fusion genes and trans-splicing events, ChimPipe is able to detect read-through events, which is now recognized as the most prevalent class of real chimeras in both normal and tumour tissues [20, 29, 30]. ChimPipe is general enough to be able to work on any eukaryotic species with a genome and an annotation available. This allows to study chimera evolution but also to investigate the impact of chimeras on individuals from species on which we have more control than human (for example livestock). ChimPipe can also predict several isoforms per gene pair and the exact chimeric junction coordinates, which are essential for chimeric transcript reconstruction and downstream validation.

ChimPipe is easy to run since it only requires a genome, a gene annotation and RNA-seq fastq files (once the indexing of the genome and transcriptome has been done), and guesses many other things such as the directionality, the mate configuration and the Illumina offset quality. For advanced users, many parameters, such as expression threshold or parent gene sequence similarity threshold, can be tuned. ChimPipe provides both a complete and a filtered set of chimeric junctions, with additional information about them, such as chimera category, expression support and the list of reads supporting the junction (Additional data file 1). In addition to chimeric junctions, ChimPipe provides a standard bam file obtained from the GEMtools RNA pipeline (step (i) of Figure 2), that can be used for downstream analyses such as differential gene expression or transcript reconstruction.

Benchmarking of ChimPipe together with four state-of-the art chimera detection tools on both simulated and real data, showed ChimPipe to have a very good precision (close to 1), and to be second most sensitive program (Sn of 0.8), therefore showing a very good balance between sensitivity and precision. ChimPipe’s performances on simulated and real data are comparable, and not much impacted by read length. They are also similar at the gene pair and the junction level, which is not the case for all programs since they generally predict gene pairs better than junctions. It has to be noted that ChimPipe needs non negligible computer resources to achieve these results, since it requires 30Gb of RAM and half a day to run with 4 threads, on the PE76 simulated data (21 million PE reads).

The application of ChimPipe to 106 ENCODE PE RNA-seq samples allowed the detection of 137 highly reliable chimeras, of which 4 were chosen for RT-PCR validation, and of which 3 were indeed validated and further cloned and sequenced. The UBA2-WTIP chimera additionally preserved the frame of the two parent genes UBA2 and WTIP, and was therefore completely sequenced. This gave rise to 3 novel transcripts which if translated, would lead to a chimeric protein with the ThiF and the UAE-Ubf domains from the UBA2 protein and with the 3 LIM domains from the WTIP protein. We hypothesize that this protein may induce the SUMOylation machinery to post-translationally modify and regulate novel targets, due to the interaction of its LIM-domains with novel proteins.

Despite these advantages, ChimPipe could be improved in at least two aspects: (1)it could provide all the chimeric transcripts compatible with the chimeric junction (module for which we already have a tested code) as additional information, (2) it could be made more robust by being reimplemented in a pipeline specific language such as nextflow http://www.nextflow.io/.

Finally it has to be noted that our contribution goes beyond the ChimPipe program, since we provide two additional programs: (1) a chimera simulator program, called ChimSim (https://github.com/Chimera-tools/ChimSim), and (2) a chimera benchmark program, called ChimBench (https://github.com/Chimera-tools/ChimBench). We also provide new realistic simulated data, as well as junction coordinates for validated fusion genes from two extensively used gold-standard chimera datasets [25, 45]. We think that, in addition to ChimPipe, both these programs and this data can be very useful in future chimera detection assessments.

## List of abbreviations

- EST =: Expressed Sequence Tag
- cDNA =: complementary DNA
- mRNA =: messenger RNA
- 3D =: three dimensional
- RT-PCR =: reverse transcriptase polymerase chain reaction
- PE =: paired end
-bp =: base pair
-kb =: kilobase
-RAM =: random access memory
-Gb =: Gigabyte

## Data availability

The ChimPipe program is available at https://github.com/Chimera-tools/ChimPipe, its associated documentation at https://chimpipe.readthedocs.io/en/latest/, and a description of its main output in Additional data file 1. The chimera simulation data is provided in Additional data file 2 at http://public-docs.crg.es/rguigo/Papers/ChimPipe/Paper/additional.data_2.tar.gz. The exact junction coordinates for the Berger and Edgren validated gene fusions are provided in Table S1. The RT-PCR validation data are provided in Additional data file 3 at http://public-docs.crg.es/rguigo/Papers/ChimPipe/Paper/additional.data_3.tar.gz. The ChimSim chimera simulation program is available at https://github.com/Chimera-tools/ChimSim, and the ChimBench chimera detection benchmark program is available at https://github.com/Chimera-tools/ChimBench.

## Ethical approval

No ethical approval was required for this study.

## Authors’ contributions

BRM wrote the ChimPipe program from an initial version from SD and EP. BRM and SD conducted the bioinformatics analyses. SM, TG and PR developed the GEMtools and the GEM RNA mapper underlying ChimPipe. GA, AR and BA performed the experimental validation (RT-PCR, cloning and sequencing). RG provided useful comments about the chimpipe method and the biological results. SD supervised the project. SD and BR wrote the manuscript with help from BA, AR, PR and RG . All authors read and approved the manuscript.

## Acknowledgements

We would like to thank Thomas Gingeras for insightful comments about ChimPipe results, Vincent Lacroix for initial discussion about chimera mechanisms, David Torrents for critical comments about benchmarking, and Rory Johnson for fruitful discussion about chimera validation. We would also like to thank Carmen Arnan Ros for technical support in the experimental validation, and Rory Johnson and Kylie Munyard for reviewing this manuscript.

## Supplementary Figures

**Figure S1.**
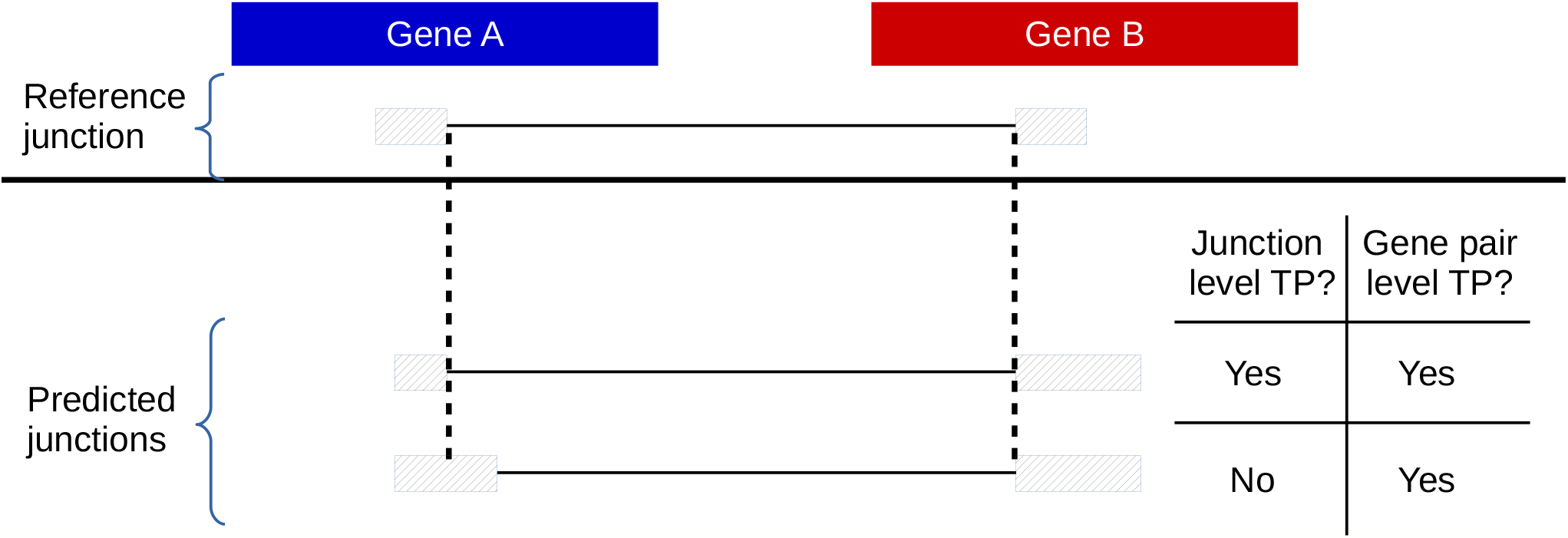
Two evaluation levels: gene pair level and junction level. On top is one reference junction with its associated gene pair (gene A, gene B). At the bottom are two predicted junctions of which the first one exactly corresponds to the reference junction, and is therefore considered both a junction level and a gene pair level true positive (TP). The second junction does not exactly correspond to the reference junction, and is therefore not considered a junction level TP, however since its first part overlaps an exon of gene A and its second part an exon of gene B, it is considered a gene pair level TP.

**Figure S2.**
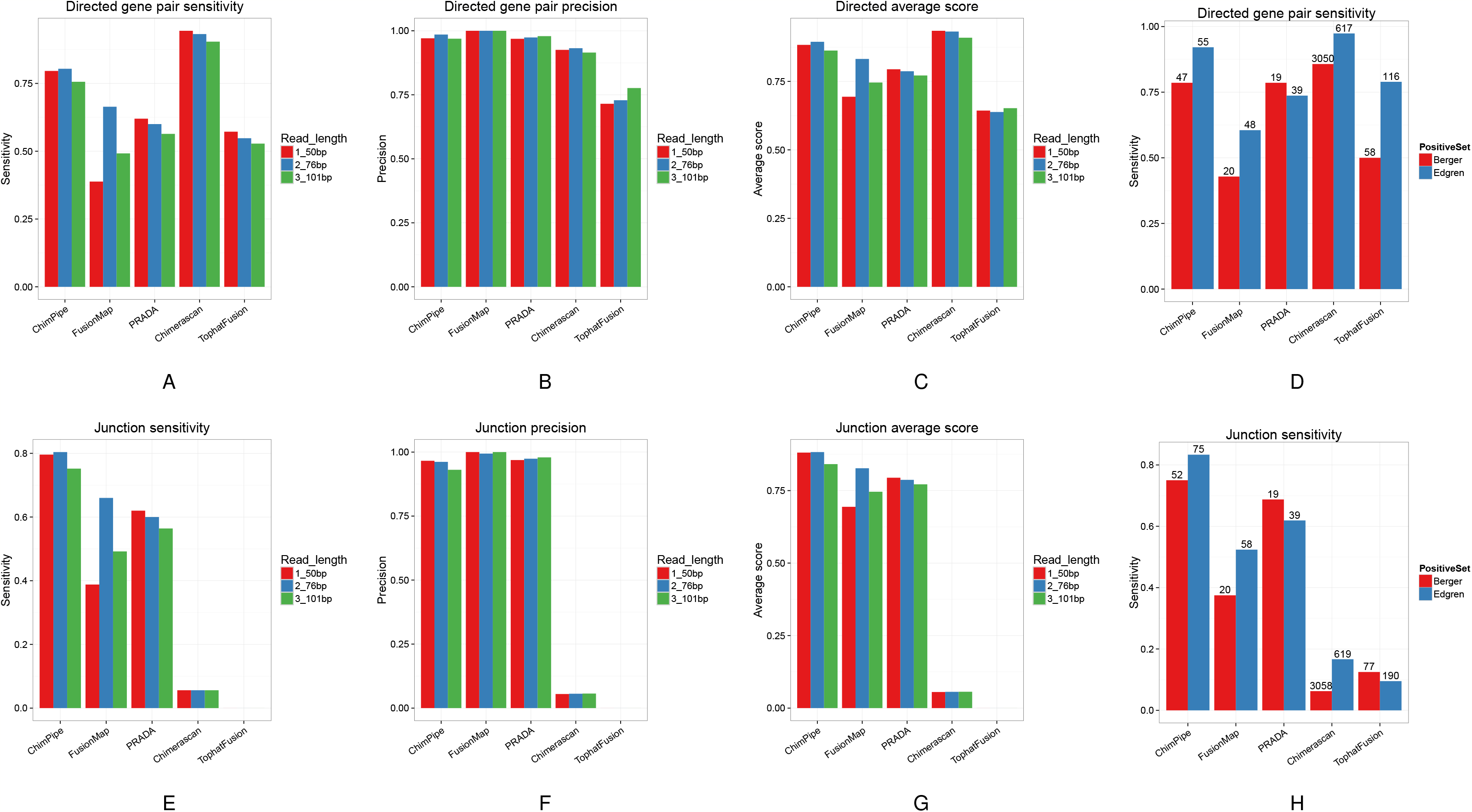
Benchmark results of 5 chimera detection programs on simulated data of different read lengths and on real data. On the top row (A-D) are the programs’ performances at the gene pair level, and on the bottom row (E-H) at the junction level. The three first sets of barplots of each row are the results on simulated data, with sensitivity in first column, precision in second column and the average between the two in third column, with different colors for 3 read lengths, while the last set of barplots of each row show the results on real data (Berger in red and Edgren in blue). According to simulated data, read length does not have a big impact on the results, except for Fusionmap which is better with 76bp reads. The sensitivity tends to decrease when the read length increases

**Figure S3.**
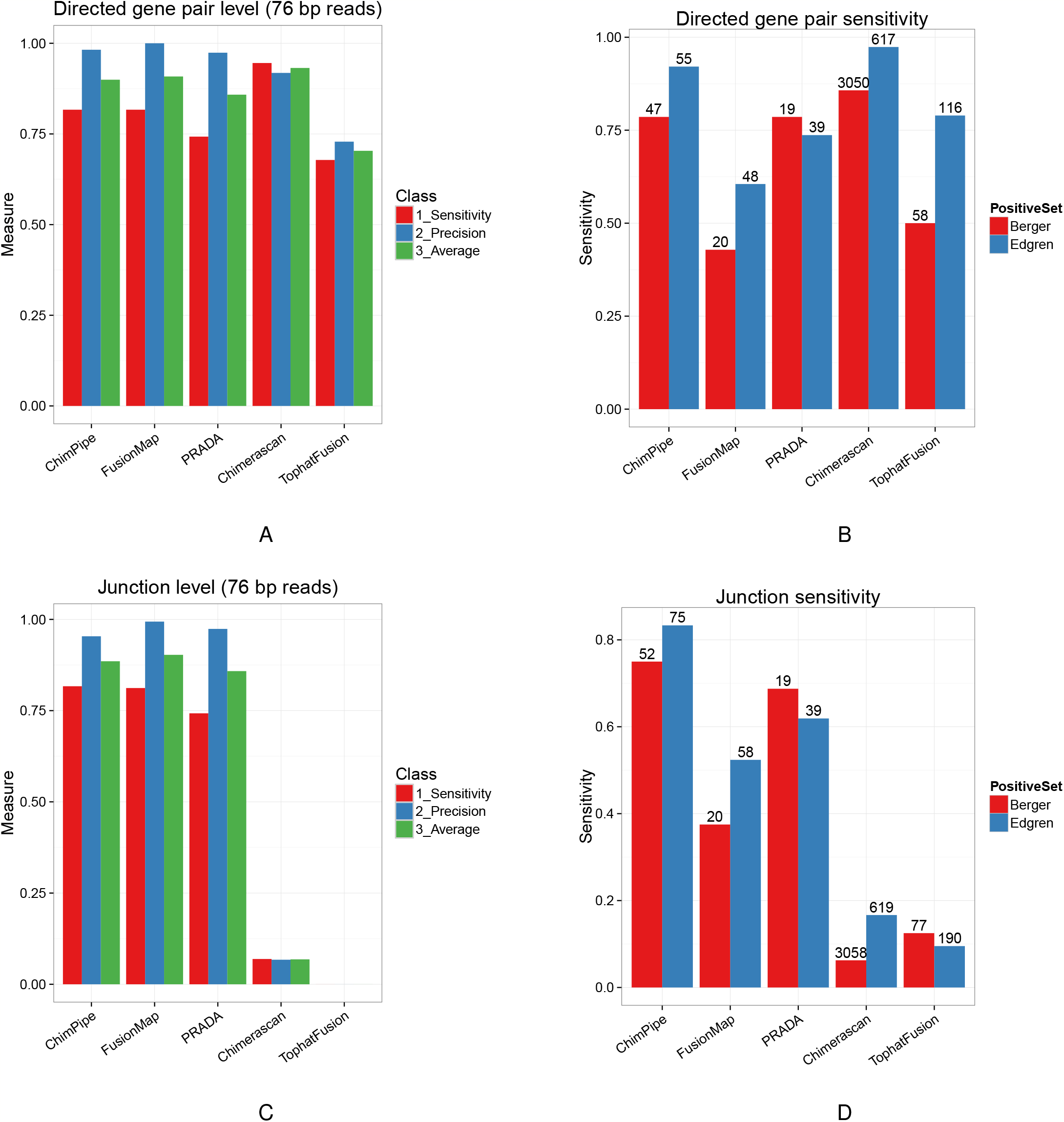
Benchmark results when excluding read-through events for simulated data. This is the same as figure 3 but excluding read-through events when benchmarking the programs on simulated data. Note that figures S3B and S3D are the same as figures 3B and 3D, and are present here for the purpose of comparison between results on simulated and real data.

**Figure S4.**
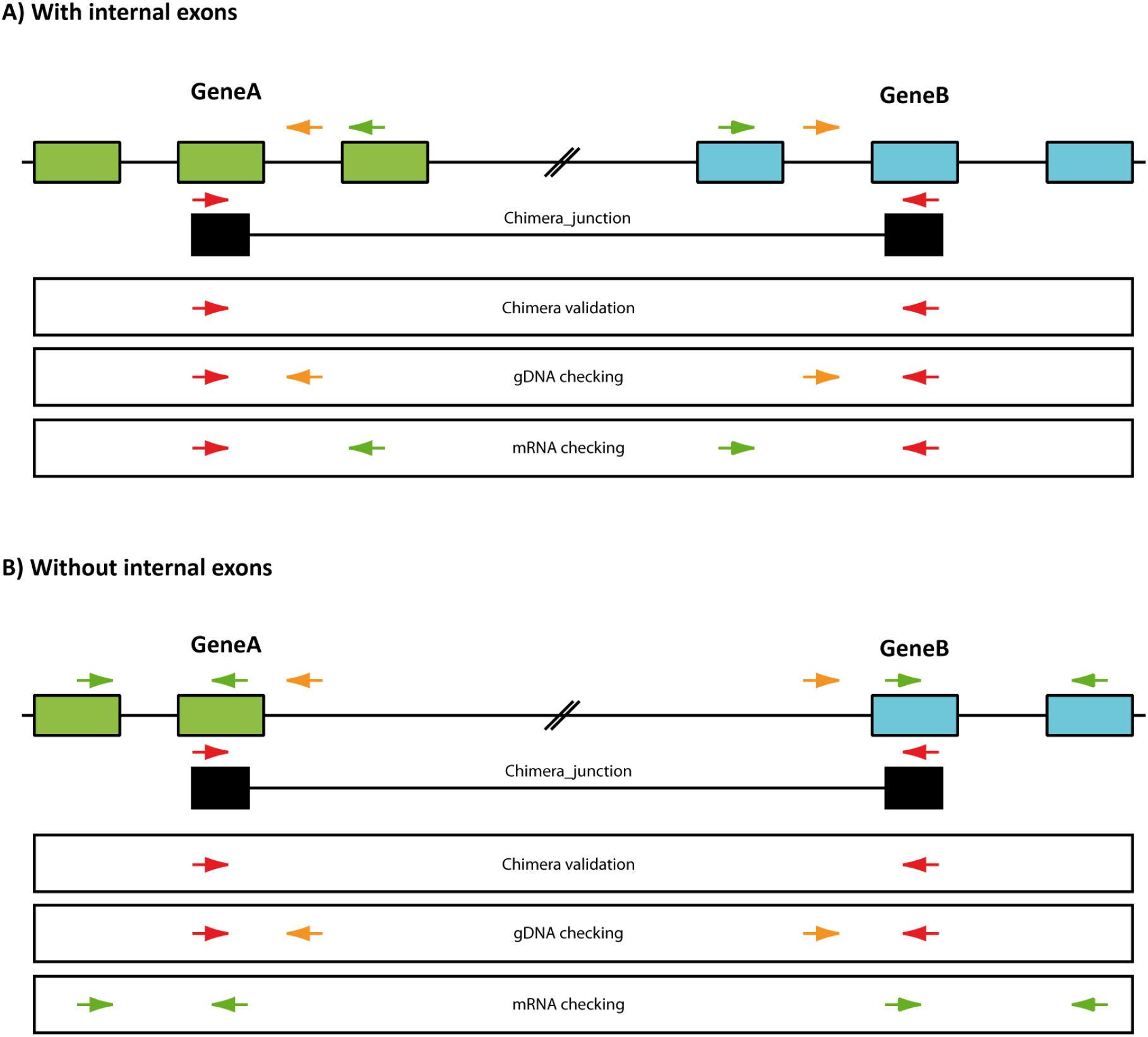
RT-PCR validation method. For each chimeric junction attempted to be validated by RT-PCR, 3 tests were actually performed, each of them requiring different pairs of primers: (1) the actual validation of the chimeric junction is done by doing RT-PCR on a cDNA library using a pair of primers that are located externally to the junction but in the exons overlapped by each part of the junction; (2) genomic DNA tests (starting from genomic DNA) are done for each of the parent gene and for the chimeric junction, in order to check whether a genomic rearrangement could explain the chimeric junction and to see if the parent genes are present at the DNA level; (3) mRNA checks are done for each parent gene (starting from cDNA and using pairs of primers specific to each gene) in order to check whether they are present at the mRNA level.

**Figure S5.**
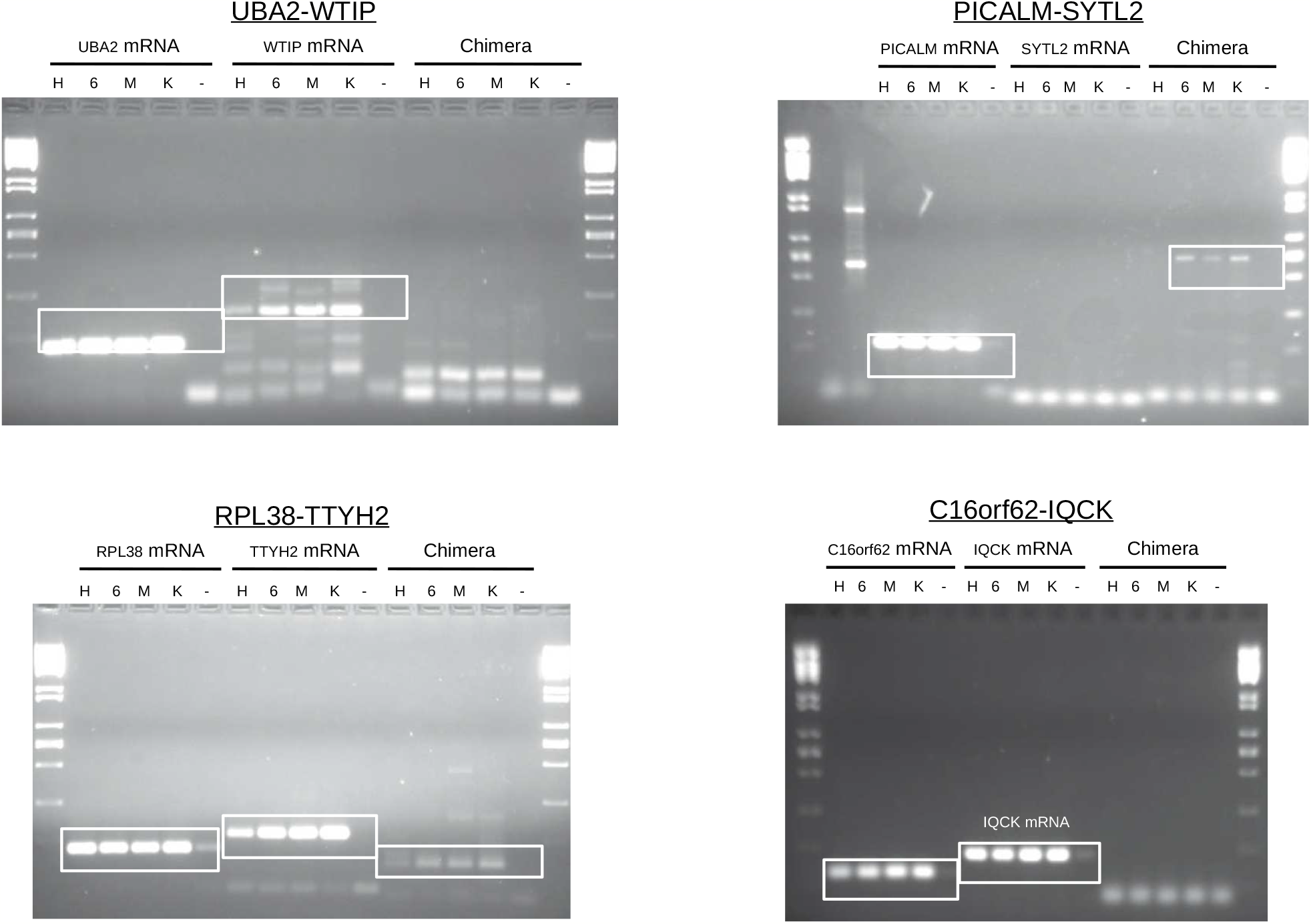
RT-PCR validation results for 4 chimeras in four cell lines (HeLa, HL60, MCF-7, K562). Here we show the products of the RT-PCR amplification of 4 chimeras and their parent genes, from the cDNAs of 4 different cell lines: HeLa (H), HL60 (6), MCF-7 (M) and K562 (K). For each chimeric junction we also provide a negative control (-) for comparison, and higlight the bands that show the presence of the chimeras and of the parent genes. For the 3 successfully validated cases (3 first ones, i.e. UBA2-WtIp, PICALM-SYTL2 and RPL38-TTyH2), the bands show the presence of the chimeric RNAs at the expected size, except for PICALM-SYTL2, which chimera size is higher than expected, and of all parent mRNAs except SYTL2, although this could be due to a low expression level of this gene.

**Figure S6.**
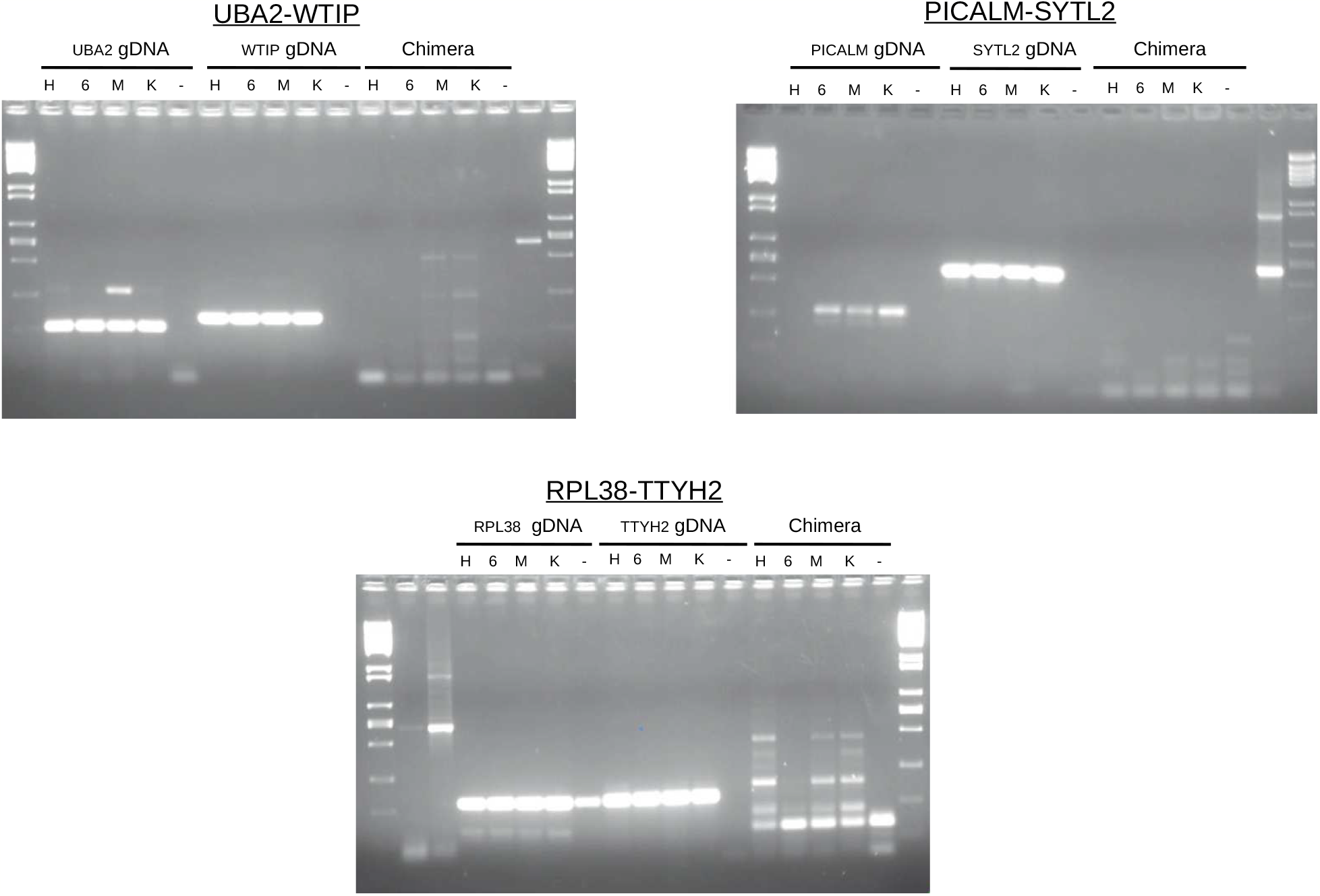
Genomic DNA analysis for the 3 successfully validated chimeras. For the 3 chimeras that were successfully validated by RT-PCR, we show the products of the genomic amplification of the 6 chimeras and their parent genes in the same 4 cell lines as the ones where the mRNA analysis was done (see supplementary Figure S5 above). These tests show that the parent genes are present at the DNA level, but not the chimeras. Indeed we see some unspecific amplification in the genomic DNA for the chimeras, but the band intensities are too low to consider them as genomic rearrangements. For the RPL38-TTYH2 chimera, there are some clearer unspecific products, but they are probably due to a primer contamination since they are also present in the negative control. H: HeLa, 6: HL60, M: MCF-7, K: K562, -: negative control.

**Figure S7.**
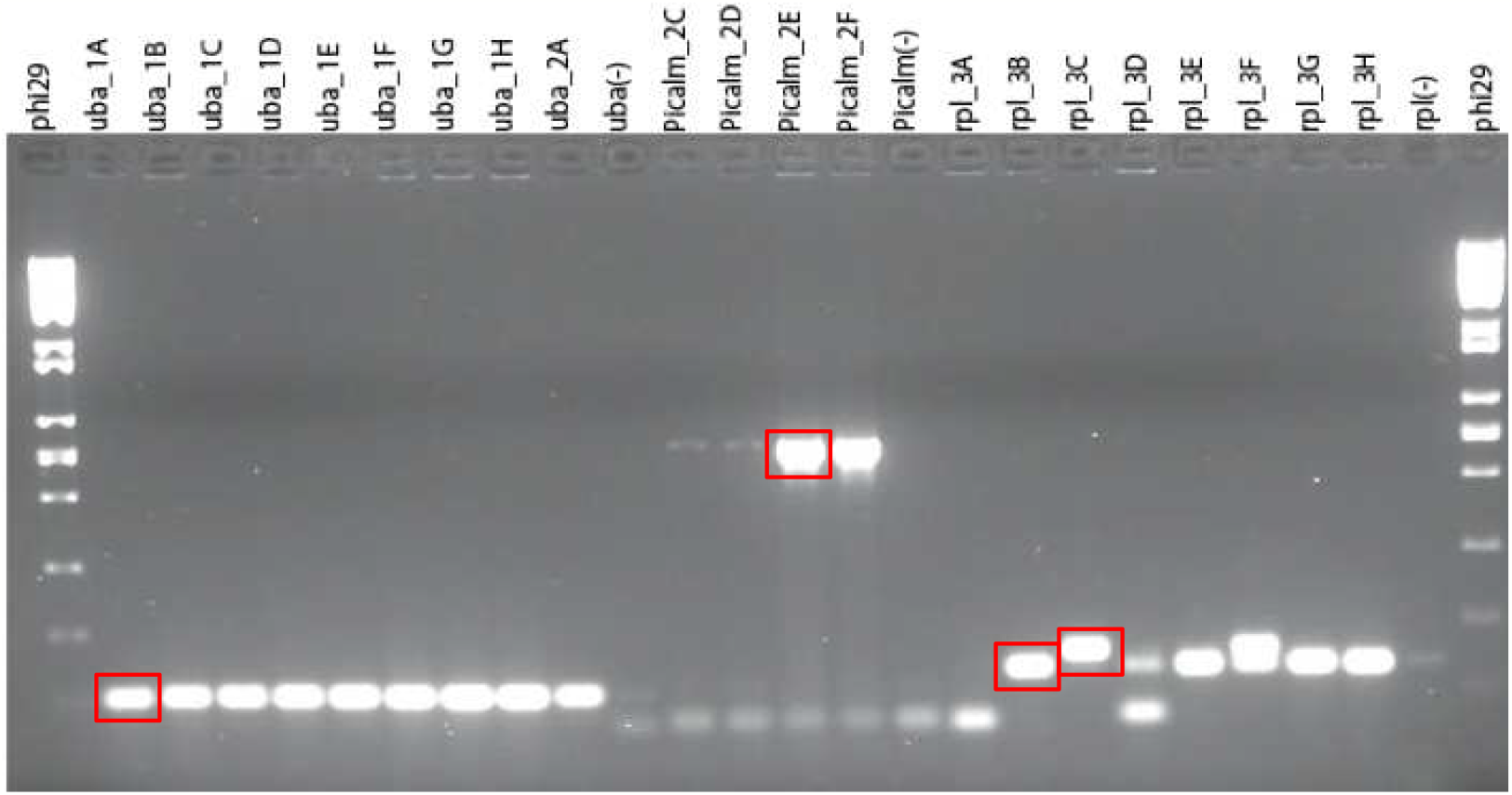
Colony PCR check and selection for RT-PCR validated chimeras. For the 3 RT-PCR validated chimeras, the RT-PCR products were purified from the gel bands, and cloned into pGEMTeasy vectors. E. coli bacterias were then transformed with these vectors, white colonies were selected and colony pCr was performed with the results indicated on the picture. Selected colonies (indicated by red boxes) were grown, plasmid purified and sent for sequencing.

**Table S1.**
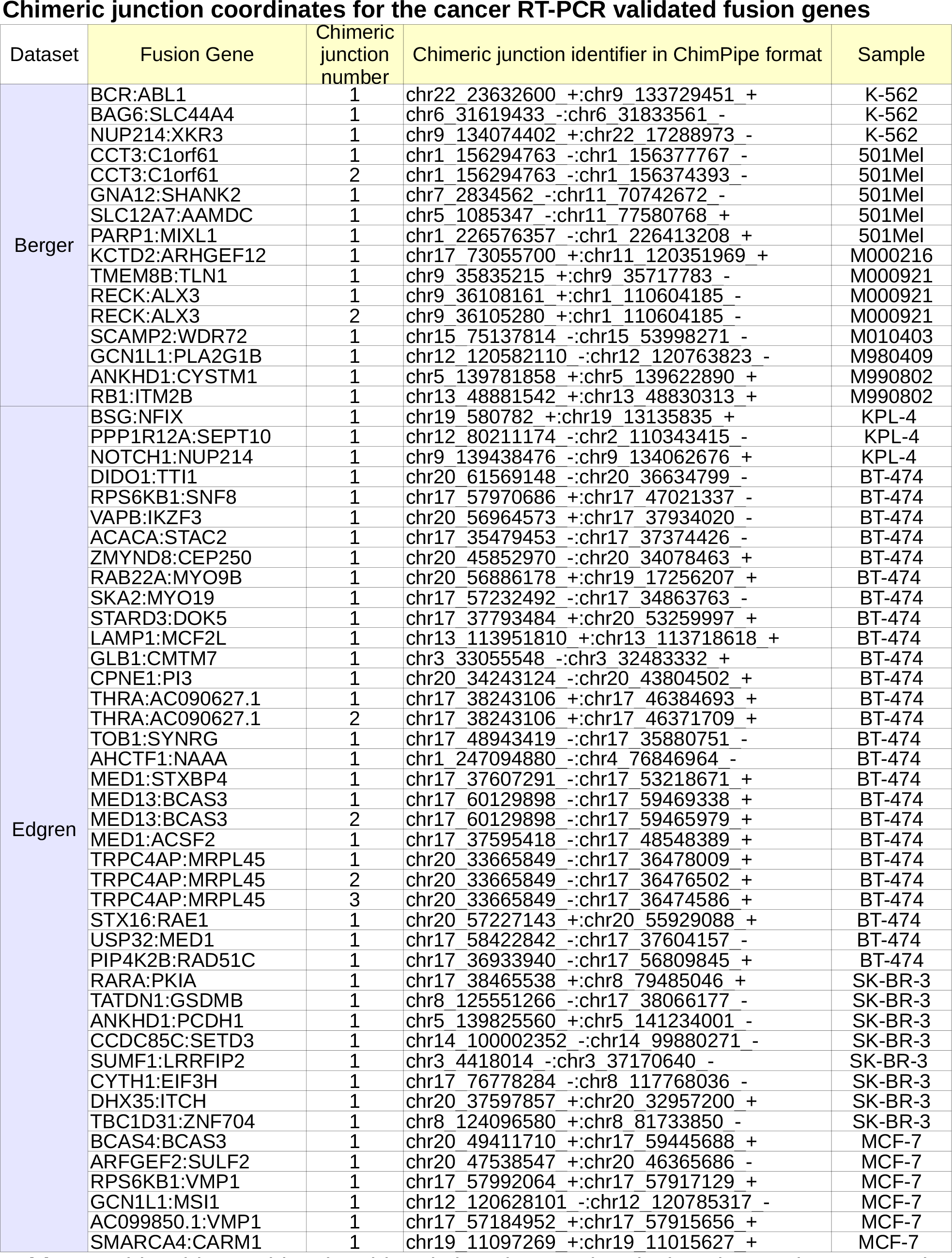
This table provides the chimeric junction number, its location on the genome in chimpipe format and the sample where it was validated, for each validated fusion gene in each of the two cancer datasets

